# Comparative transcriptomics reveals divergence in pathogen response gene families amongst 20 forest tree species

**DOI:** 10.1101/2023.03.06.531373

**Authors:** Mengmeng Lu, Min Cao, Jie Yang, Nathan G. Swenson

**Author notes:** Corresponding author: Department of Biological Sciences, University of Notre Dame, 100 Galvin Life Sciences, Notre Dame, IN 46556, USA.

## Abstract

Forest trees provide critical ecosystem services for humanity that are under threat due to ongoing global change. Measuring and characterizing genetic diversity is key to understanding adaptive potential and developing strategies to mitigate negative consequences arising from climate change. In the area of forest genetic diversity, genetic divergence caused by large-scale changes at the chromosomal level has been largely understudied. In this study, we used the RNA-seq data of twenty co-occurring forest trees species from genera including *Acer*, *Alnus*, *Amelanchier*, *Betula*, *Cornus*, *Corylus*, *Dirca*, *Fraxinus*, *Ostrya*, *Populus*, *Prunus*, *Quercus*, *Ribes*, *Tilia*, and *Ulmus* sampled from Upper Peninsula of Michigan. These data were used to infer the origin and maintenance of gene family variation, species divergence time, as well as gene family expansion and contraction. We identified a signal of common whole genome duplication events shared by core eudicots. We also found rapid evolution, namely fast expansion or fast contraction of gene families, in plant-pathogen interaction genes amongst the studied diploid species. Finally, the results lay the foundation for further research on the genetic diversity and adaptive capacity of forest trees, which will inform forest management and conservation policies.

## Introduction

Forest trees occupy nearly a third of the Earth’s land surface and provide key ecological benefits and services such as water, air purification and carbon sequestration (FAO 2020; Gamfeldt et al. 2013; Miura et al. 2015). They are critical components of terrestrial biodiversity in terms of their own diversity, but also with respect to the habitats and renewable materials such as wood, cellulose, and lignin (Cortés et al. 2020). It is estimated that there are ∼73,000 tree species globally with most of being rare and regionally endemic, implying vulnerability of global tree species to anthropogenic and climate changes (Cazzolla Gatti et al. 2022). Indeed, climate change has already impacted forests and their species composition via range migration, wildfires, droughts, pest, and pathogens (Allen et al. 2010; Hammond et al. 2022; Kirilenko and Sedjo 2007; Rizvi et al. 2015). The genetic composition and diversity of the tree species in forests is foundational not only to our understanding of the abiotic and biotic factors governing the distribution and dynamics of species populations, but also to how they will respond to current and future environmental change. Thus, measuring and characterizing intra-and inter-specific genetic diversity is the key for understanding the current structure and dynamics of forests and their adaptive potential and for developing strategies to mitigate the damages brought along with environmental change.

Gene duplication and gene loss are major sources of genetic diversity. Gene duplication, generated through either whole genome duplication (WGD) or small-scale duplication (SSD), is the primary source of new genetic material among flowering plants (Cui et al. 2006; Panchy et al. 2016). WGD contributes to the majority of duplicate genes (Amborella Genome Project 2013; Soltis and Soltis 2016). The ancient WGDs can lead to competitive advantage, increased species diversity, and evolutionary innovations (Van de Peer et al. 2009). Most flowering species have a polyploid history either through autopolyploidy or allopolyploidy. Specifically, flowering species are either currently polyploid, or have a polyploid ancestor and have reverted to being diploid (Blanc and Wolfe 2004). Earlier studies placed at least three WGD events during the radiation of angiosperms, the oldest WGD occurred before the monocot-eudicot divergence, a second WGD occurred specifically to eudicots, a more recent WGD, gamma (γ) triplication event, occurred before the separation of rosids and asterids (Jiao et al. 2012; Tang et al. 2008). Alternatively, pervasive SSD, including tandem duplication (Zhang 2003), transposon-mediated duplication (Jiang et al. 2004), segmental duplication (Bailey et al. 2002), and retroduplication (Brosius 1991), also generates a great number of duplicate genes. While the majority of duplicate genes end up being silenced and eliminated (Lynch and Conery 2000), a few preserved duplicate genes may be retained by selection due to neo-functionalization, relative dosage, absolute dosage or sub-functionalization (Conant et al. 2014), and play important role in generating genetic diversity and building up reproductive isolation barriers. Recent studies explained retention of duplicate genes as some paralogous genes depend on or compensate for their cognate copy (Diss et al. 2017), while some paralogous gene pairs are stuck in a functional and structural entanglement status over evolutionary time scale (Kuzmin et al. 2020).

Compared to gene duplication, the mechanism that underlie gene loss are not yet well understood. It is commonly assumed that gene loss is realized by deletion or pseudogenization, as a result of neutral or adaptive processes (Demuth and Hahn 2009). Most nonsense mutations are subject to drift to fixation. Some deleterious mutations could be accumulated and eventually become silenced (Johri et al. 2022). Gene loss might be selected to retain dosage balance, as evidenced in a set of single-copy genes with essential housekeeping functions, which are typically restored to become singletons after WGD or SSD (De Smet et al. 2013).

Differential duplication and loss of genes among evolutionary lineages results in gene family expansion and contraction (Demuth and Hahn 2009). Gene families comprise a set of similar nucleotide or amino acid sequences. They arise as a result of duplication events and have similar cellular and biochemical functions. Gene family expansions and contractions related to metabolic, regulatory and signalling gene networks contribute to the adaptations of plants to various abiotic habitats and biotic interactions (One Thousand Plant Transcriptomes Initiative 2019). They are sources of plant functional evolution and innovation (Force et al. 1999; Ohno 1970), exhibiting a clear functional basis, which are influenced by the mechanisms of duplication (Hanada et al. 2008; Maere et al. 2005). Comparisons of gene family content across species, therefore, may provide insight into the ecological interactions and evolutionary pressures that have shaped adaptation and diversity. This is particularly true when we conduct comparative transcriptomic analyses of species that currently co-occur and that, potentially have co-occurred for long periods of time (Swenson and Jones 2017).

Genetic investigations of forest trees have, generally, focused on nucleotide substitutions in specific genes (Isabel et al. 2020; Neale and Kremer 2011), while comparative analyses of genetic divergence caused by large-scale changes at the chromosomal levels have been largely understudied. However, the magnitude, rate, and distribution of changes in gene family size are key drivers of the functional differentiation and adaptation of species (Demuth and Hahn 2009; Lynch and Conery 2003). Large segmental duplication and changes in copy number are known to be associated with disease response and environmental adaptation (Bailey et al. 2002; Reid et al. 2016). Therefore, gene family size change is as much important as nucleotide substitution to understanding genetic divergence and evolutionary forces responsible for these changes.

Such knowledge on genetic divergence and associated evolutionary forces in forest trees can help to promote breeding healthy trees (i.e. disease resistant trees) in a changing climate (Allen et al. 2010; Dodds and Rathjen 2010). The increasing occurrences and severity of insect and disease outbreaks have threatened forest trees (Hurley et al. 2016; Linnakoski and Forbes 2019), but we lack a thorough understanding to the disease related genes in forest trees. Trouern-Trend et al. (2020) revealed contraction in disease associated gene families in *Juglans* species. Lu et al. (2021) reported differential gene expression patterns between tolerant and susceptible lodgepole pine trees to a fungal pathogen. It would be interesting to explore the potential link between duplication events and disease response in expressed genes of forest trees, as the duplicate genes can quickly lead to changes in gene expression with the acquisition of increased resistance to different pathogens (Picart-Picolo et al. 2020).

In this study, we used the RNA-seq data for twenty co-occurring forest trees species to infer the origin and maintenance of gene family variation as well as species divergence time, gene expansion and contraction. We sought to address the following questions: What is the magnitude of gene duplication? What factors impact duplicate gene evolution? Are the gene gain and loss process random or controlled by natural selection? The answers to these questions are the key to understanding the basis of adaptive divergence in forest tree species and will help to predict adaptive capacity of these species in a changing climate.

## Materials and methods

### Sample collection

Leaves of each of twenty forest species were sampled in August, 2021, at the University of Notre Dame Environmental Research Center (UNDERC) near Land O’ Lakes, Wisconsin, U.S.A. (Fig. 1). The UNDERC site is located at latitude: 46.23391, longitude: -89.537254, and encompasses approximately ∼8,000 acres of mixed deciduous forest on both sides of the state line between Wisconsin and Michigan’s upper peninsula. The twenty species used in this study are dominate components of the woody vegetation at the site and are: *Acer rubrum* (red maple), *Acer saccharum* (sugar maple), *Acer spicatum* (mountain maple), *Alnus incana* (Gary alder), *Amelanchier laevis* (smooth serviceberry), *Betula alleghaniensis* (yellow birch), *Betula papyrifera* (paper birch), *Cornus alternifolia* (pagoda dogwood), *Corylus cornuta* (beaked hazelnut), *Dirca palustris* (Eastern leatherwood), *Fraxinus nigra* (black ash), *Ostrya virginiana* (American hophornbeam), *Populus grandidentata* (bigtooth aspen), *Populus tremuloides* (quaking aspen), *Prunus serotina* (black cherry), *Prunus virginiana* (chokecherry), *Quercus rubra* (northern red oak), *Ribes cynosbati* (prickly gooseberry), *Tilia americana* (American basswood), *Ulmus americana* (American elm). They are all angiosperms (flowering plants) and eudicots, belonging to the clades of asterids, saxifragales, and rosids. These species were chosen as they are the most dominant at the study site and in large national forests in region (i.e. Ottawa, Nicolet, and Hiawatha National Forests). Specifically, these species compose >95% of the tree individuals and above-ground biomass in these forests (Swenson et al. 2017). We focus on the most dominant species in order to provide insights into the ecological and evolutionary processes driving transcriptomic variation within and across populations and communities (Swenson and Jones 2017). Upon collection, the samples were flash frozen in liquid nitrogen, then transported and stored in an ultracold freezer at the field station.

**Fig. 1.**
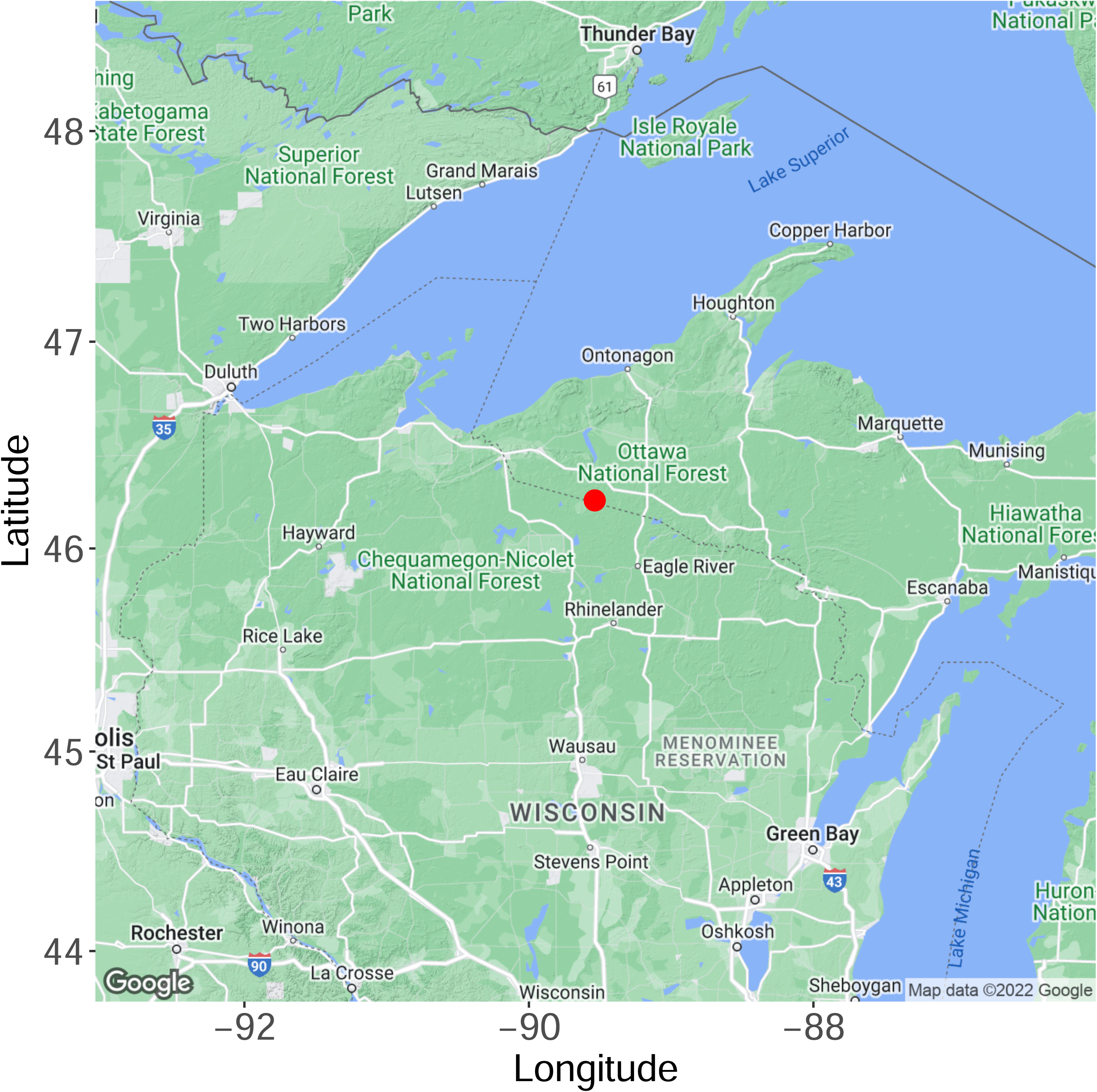
The location (dot) of the University of Notre Dame Environmental Research Center (UNDERC) where all samples used in this study were collected.

### RNA-sequencing, de novo transcriptome assembly, and annotation

The samples were sent to Novogene corporation inc. (Chula Vista, CA, USA) for mRNA extraction, library preparation, mRNA sequencing, and quality control. The samples were sequenced on an Illumina NovaSeq 6000 platform to produce 150 bp paired-end reads. Raw reads filtering was performed by Novogene to get the clean reads including removing reads containing adapters, removing reads containing N >10%, and removing reads containing low quality base (Phred quality score ≤5) which is over 50% of the total base. An average of 6 Gb paired-end clean reads were acquired for each sample and deposited in NCBI Sequence Read Archive (SRA accession number: PRJNA896247). *De novo* transcriptome assembly was performed for each sample using the software TransLiG v1.3 with default parameters (Liu et al. 2019). The clean reads were then mapped to the assembled transcriptome using the software RSEM v1.2.25 with default parameters (Li and Dewey 2011). The longest transcript (isoform) with a transcript per million (TPM) value ≥1.5 for each gene was retained. Redundancy was removed from the retained transcripts using the software CD-HIT v4.8.1 with parameters of -c 0.95 -n 8 -T 32 -M 1540 (Fu et al. 2012). To remove potential contamination, the retained transcripts were aligned against the non-redundant (nr) protein database (downloaded on January 22, 2022) using the blastx function implemented by the software DIAMOND v2.0.13.151 (Buchfink et al. 2021) with parameter of --outfmt 102 to assign the best known taxon ID to each query transcript. The file that was used to convert accession numbers to taxonomy was downloaded from https://ftp.ncbi.nlm.nih.gov/pub/taxonomy/accession2taxid/, and the taxon ID file was downloaded from https://ftp.ncbi.nlm.nih.gov/pub/taxonomy/new_taxdump/. The query transcripts that could be assigned to green plant (Viridiplantae) taxon IDs were retained. The filtered transcripts were aligned to the nr database that only includes green plant subjects using the blastx function implemented by the software DIAMOND v2.0.13.151 with default parameters. The aligned transcripts were combined with those transcripts assigned with green plant taxon IDs to form the final transcriptomes. The completeness of the final transcriptomes was examined using the 1,614 orthologues in the BUSCO (Benchmarking Universal Single Copy Orthologs) v5.2.2 set of embryophyta_odb10 (Manni et al. 2021).

The transcriptomes were annotated with Gene Ontology (GO) and Kyoto Encyclopedia of Genes and Genomes (KEGG) terms. The transcripts were aligned to the nr database that only includes green plant subjects using the blastx function implemented by the software DIAMOND v2.0.13.151 with parameters of -taxonlist 33090 --very-sensitive -f5. The .xml outputs were used as input for mapping and annotation using the software Blast2GO (Götz et al. 2008). The transcripts were translated to protein sequences using the software TransDecoder v5.5.0 (https://github.com/TransDecoder/TransDecoder/wiki) with default parameters to get the longest open reading frame (ORF) per transcript, and then were annotated by the software InterProScan v5.54-87.0 (Jones et al. 2014) with the parameters of -dra -f XML -goterms -pa. The final GO terms were acquired by merging Blast2GO and InterProScan annotation outputs using the software Blast2GO. To get the KEGG terms, the longest ORF per transcript were also annotated using EggNOG-mapper v2.1.7 (Cantalapiedra et al. 2021) with parameters of -m diamond - sensmode more-sensitive -tax_scope 33090 -tax_scope_mode 33090. The assembled transcriptome and annotation were used in the downstream analyses as shown in the flowchart (Supplementary Fig. S1).

### Ks analyses

The level of synonymous substitutions (Ks) between two homologous sequences increases approximately linearly with time. Thus pairs of paralogous genes within a species can be sorted to estimate the relative ages of gene duplication (Blanc and Wolfe 2004). To identify paralogous gene pairs within a species, an all-versus-all blastp was performed within each species. The peptide sequences (the longest ORF per transcript) of each species were used as input to run all-versus-all blastp using the software DIAMOND v2.0.13.151 with parameters of -p 16 --max-target-seqs 20 --evalue 1e-10 --outfmt 6 qseqid sseqid pident nident evalue. The blastp output was filtered by removing self blastp results, keeping outputs with pident (percent of identical matches) ≥20 and nident (number of identical matches) ≥50, and removing sequences with more than 10 hits. The filtered paralogous gene pairs were aligned using the software MAFFT v7.490 (Katoh et al. 2002) with default parameters. The alignments were used as input for estimating the number of synonymous substitutions per synonymous site (Ks) and the number of non-synonymous substitutions per nonsynonymous site (Ka) using the program yn00 in the software PAML v4.9j (Yang 2007) with parameters of icode = 0 weighting = 0 commonf3×4 = 0. The Ks of each paralogous gene pairs were acquired using the method by Yang and Nielsen (2000).

Ks can also be calculated using the orthologous gene pairs between species to estimate speciation events (Blanc and Wolfe 2004). To identify orthologous gene pairs between species, reciprocal best hit blastp were run between the compared pairs. The peptide sequences (the longest ORF per transcript) of the paired species were used as input to run blastp using the software DIAMOND v2.0.13.151 with parameters of -p 16 --max-target-seqs 20 --evalue 1e-10 --outfmt 6 qseqid sseqid pident nident evalue. The best hits between forward and reverse blastp for each compared sequence pairs were retained, which were then used for sequence alignment and estimating Ka and Ks as abovementioned.

The Ks distribution is proportional to the time lapse. Thus, the Ks distribution within species can be used to infer the timing of WGD, and the Ks distribution between two different species can infer time since speciation (Blanc and Wolfe 2004). To plot the Ks distribution, the mixture model was fit and plot for the Ks values ≥0.01 and ≤2 using the R package mixR (Yu 2022). The best model from a candidate of mixture models was selected using the select function based on the information criterion BIC.

### Estimating duplication events and divergence time

To infer duplication events, the species tree and gene trees were first inferred using STAG and STRIDE algorithms (Emms and Kelly 2017; 2018) implemented by the program OrthoFinder v2.5.4 (Emms and Kelly 2019). The inferred species tree was similar to that expected given current knowledge of the Angiosperm phylogeny with the exception of the placement of *Populus tremuloides* and *P. grandidentata*. Thus, for downstream analysis and discussion, we removed these two species for the species tree inference. The inferred species tree and gene trees were used to estimate duplication events and divergence time. The gene duplication events on the species tree node were reported if, at least 50% of the descendant species have retained both copies of the duplicate gene. The divergence time was estimated using the software r8s v1.81 (Sanderson 2003) with calibration points of 115 million years (MYA) between *Fraxinus nigra* and *Corylus cornuta*, which were obtained in November, 2022 from http://www.timetree.org/ (Kumar et al. 2017).

### Gene families and their evolutionary rate

To infer gene families, the peptide sequences (the longest ORF per transcript) of the twenty studied transcriptomes were used to construct orthogroups (gene families) using the program OrthoFinder v2.5.4 with parameters of -S diamond -t 16 -a 16. Genes assigned to each gene family were counted for each species. The evolutionary rate of protein coding genes was represented by the ratio of nonsynonymous substitutions per nonsynonymous site to synonymous substitutions per synonymous site (Ka/Ks) with *Dillenia indica* (elephant apple) being the outgroup and the twenty studied species being the ingroup. *D. indica* is a eudicot species, belonging to the order Dilleniales and the RNA-seq data for this species were downloaded from https://trace.ncbi.nlm.nih.gov/Traces/index.html?view=run_browser&acc=ERR2040190&disp lay=data-access (One Thousand Plant Transcriptomes Initiative 2019). The raw sequences were filtered using the program fastp with default parameters (Chen et al. 2018). The transcriptome assembly and protein sequence prediction were performed as abovementioned, yielding 22,753 transcripts and 19,345 protein sequences. Then the reciprocal best hit blastp was performed between *D. indica* and each of the twenty studied species to identify the orthologous pairs. The best hits between forward and reverse blastp for each compared sequence pairs were retained, which were then used for sequence alignment and estimating Ka/Ks as abovementioned. The mean Ka/Ks across genes for each gene family per species was used to represent the evolutionary rate for each gene family. Gene families with a mean Ka/Ks >10 were filtered out. The graphs were plotted using the R package ggplot2 and ggpubr (Kassambara 2023; Wickham 2016). The size and evolutionary rate of gene families were correlated using scatter plot. All statistical analyses were performed in R (R Core Team 2021). Gene families with a size ≥5 and Ka/Ks >1 were highlighted for further analysis. The enriched GO and KEGG terms were identified in these highlighted gene families. The enriched GO terms of each category (biological process, molecular function, cellular component) were separately detected using the “weight” algorithm with the Fisher ratio test implemented by the R package topGO (Alexa and Rahnenfuhrer 2023). Enriched KEGG terms were detected using the hypergeometric method implemented by the R package plyr (Wickham 2011). Heatmaps were plot using the R package pheatmap (Kolde 2019).

### Estimating gene family expansion and contraction

To further study gene family changes, species tree and gene family counts were predicted and used to estimate gene family expansion and contraction. All-versus-all blastp was performed to find the most similar sequences for each sequence in the studied species using the program DIAMOND v2.0.13.151. The program MCL (Enright et al. 2002) was used to find clusters of similar sequences (gene families). The species tree was estimated using the program FastTree (Price et al. 2009). The programs DIAMOND, MCL, and FastTree were implemented by the program OrthoFinder v2.5.4, default parameters were used. Then, the species tree was converted to a format for estimating ultrametric tree using the program CAFE5 (Mendes et al. 2021) with the command python prep_r8s.py -s 60164 -p ’Fraxinus_nigra,Corylus_cornuta’ -c 115. “60164” was the number of sites in the alignment. “-c 115” represents a median of 115 MYA divergence time, it is the calibration points between *Fraxinus nigra* and *Corylus cornuta*. Then the ultrametric tree was estimated using the program r8s v1.81 with default parameters. The ultrametric tree and gene family counts were used as input for the program CAFE5. Since CAFE5 was not recommended to analyze recent polyploid species, only diploid species were used to run CAFE5, including *Cornus alternifolia*, *Fraxinus nigra*, *Dirca palustris*, *Tilia americana*, *Acer saccharum*, *Acer spicatum*, *Amelanchier laevis*, *Quercus rubra*, *Betula papyrifera*, *Alnus incana*, *Corylus cornuta*, *Ostrya virginiana*, and *Ribes cynosbati*. Since assembly errors might cause the observed number of gene copies in gene families to deviate from the true ones and overestimation of the birth-death parameter γ (number of gene gains and losses per gene per million years), an error model was estimated first with the ultrametric tree and gene family counts using CAFE5 with parameters of -c 16 -p -e. This error model along with the ultrametric tree and gene family counts were used to estimate the birth-death parameter γ, which represents the probability that any gene will be gained or lost. Multiple γ mode was used with three different rates of gene family evolution assigned as below:

((Cornus_alternifolia:3,Fraxinus_nigra:3):3,((((Dirca_palustris:2,Tilia_americana:2):2,(Acer_sac charum:2,Acer_spicatum:2):2):2,((((Betula_papyrifera:2,Alnus_incana:2):2,(Corylus_cornuta:2, Ostrya_virginiana:2):2):2,Quercus_rubra:2):2,Amelanchier_laevis:2):2):2,Ribes_cynosbati:1):1);

The numbers 1, 2, 3 represent three different gene family evolutionary rates that were shared by different clades. This tree file was used as input along with the ultrametric tree, gene family counts, and error model using CAFE5 with the parameters of -c 16 -p -k 3. Gene families or parent nodes with a significantly greater rate of evolution (family-wide p-value <0.01), in other words, rapidly evolving gene families, were counted for each species and each parent node.

These gene families significantly violated a random gene birth and death model. Thus, the null hypothesis of neutral evolution was rejected and these families could be recognized as fast evolving gene families (Hahn et al. 2005). Enriched GO and KEGG terms were detected in the fast expanding and fast contracting gene families, respectively.

## Results

### Ks analyses

We acquired an average of 41 million raw reads for each sample. The total number of assembled transcripts ranged between 19,459 and 31,804 (Supplementary Table S1). We retrieved 76 ∼ 88% of the BUSCOs in the assembled transcriptomes (Supplementary Fig. S2). Most species have an initial Ks peak around 0.02∼0.04, except *Fraxinus nigra* and *Tilia americana*, which have the initial peak around 0.1 and 0.09, respectively (Fig. 2). Such initial peaks represent most recently duplicate genes. Another discernable separate peak shared by the studied species is around 0.5 ∼ 0.7. Such shared peaks by all species may correspond to large-scale duplication events, perhaps the common WGDs across core eudicots. Some species have other discernable peaks, like *Populus grandidentata* and *Populus tremuloides*, which have discernable peaks around 0.18. Such unique peaks may represent ancient duplication events specific to certain species and different evolutionary patterns across species. The two *Prunus* species have similar Ks shapes, so do the two *Populus* species, suggestive of common WGDs before the two species diverged from the same ancestors. By examining the orthologous Ks of the two *Prunus* and of the two *Populus* species, respectively, we found a high peak close to zero, which may represent the duplication events after the WGDs and divergence (Supplementary Fig. S3).

**Fig. 2.**
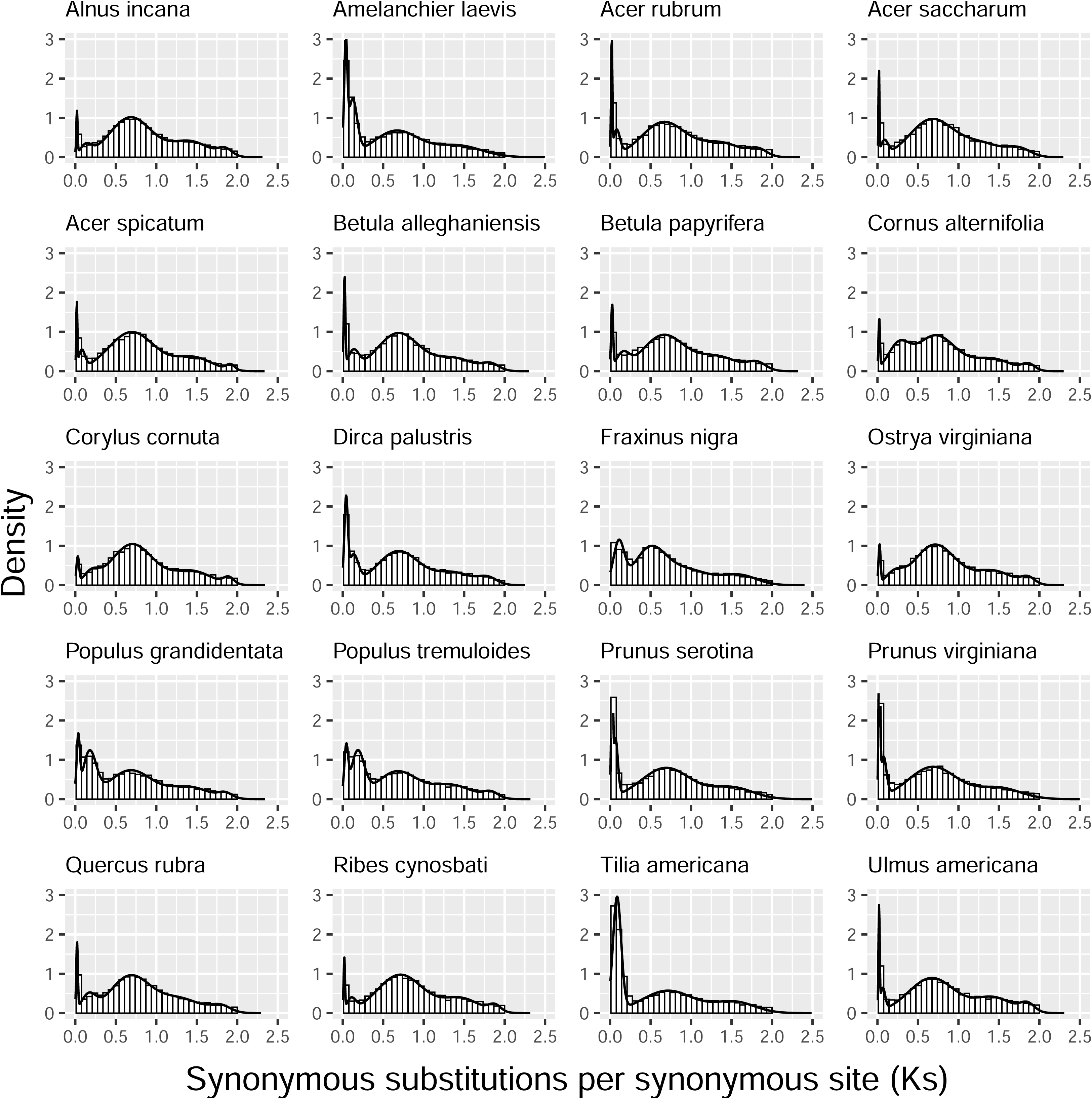
Distribution of Ks (the level of synonymous substitutions). A mixture model was fit and plot for the Ks values ≥0.01 and ≤2 of each species using the R package mixR.

### Duplication events and divergence time

Using a species tree with estimated branch lengths, we inferred divergence times and duplication events (with at least 50% support rate, Fig. 3). Most duplication events (n=1843) occurred at the divergence of rosids and asterids, which may correspond to common WGDs across eudicots species. The divergence time of the two asterids species *Cornus alternifolia* and *Fraxinus nigra* from a common ancestor was about 104 MYA (Fig. 3). The divergence time of the studied malvids and fabids from a common ancestor with Saxifragales was about 100 MYA.

**Fig. 3.**
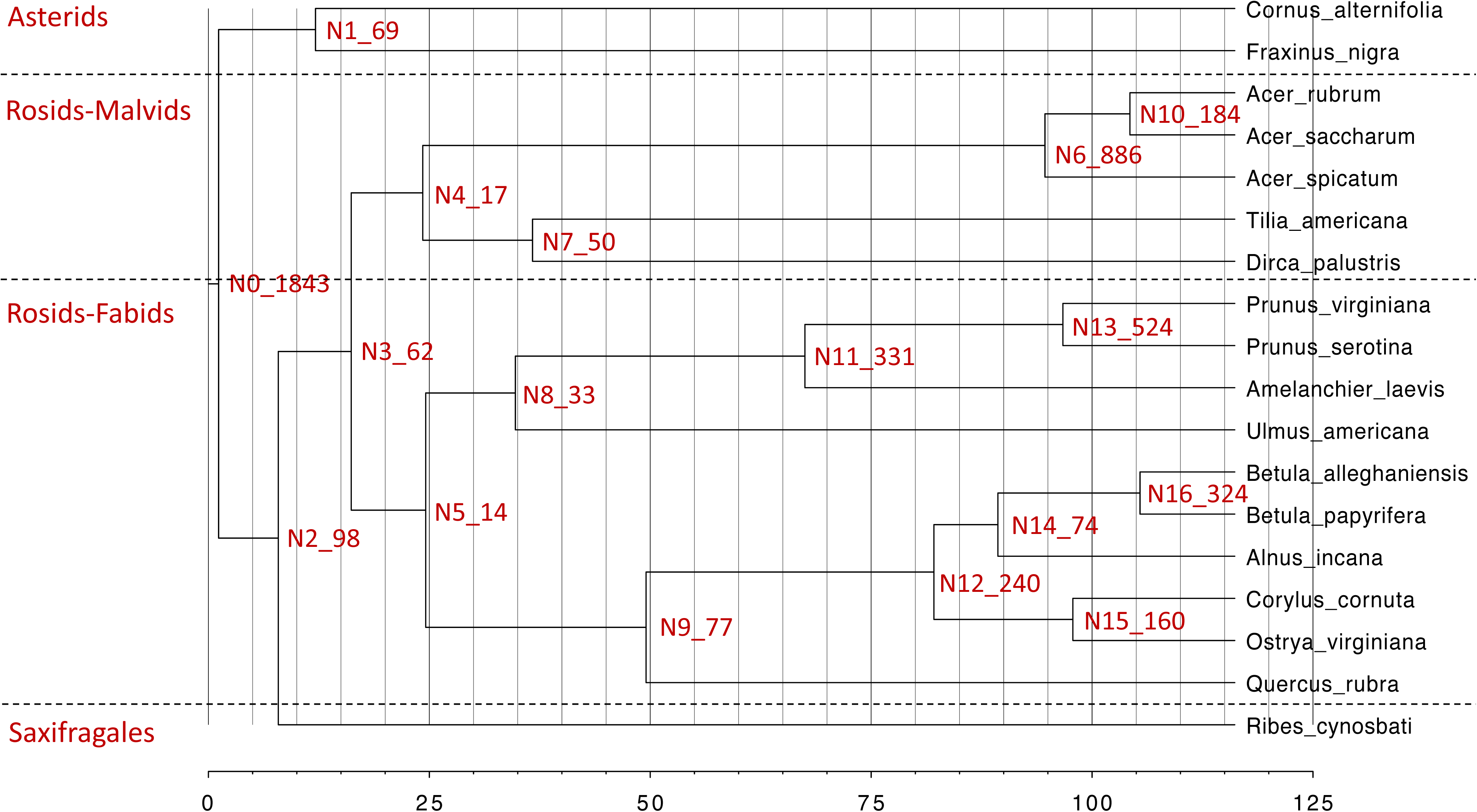
Divergence time in million years and gene duplication events with 50% support rate. Node numbers were followed by number of gene duplication events.

### Relationship between gene family size and evolutionary rate

OrthoFinder assigned 393,796 genes (96.2% of total) to 21,900 orthogroups (gene families). The mean and median gene family size is 17.3 and 12, respectively. We found that 3,060 families present in all species, whereas 19 consisted entirely of single-copy genes.

After identifying reciprocal best blastp hits with the outgroup *D. indica*, we retained a total of 6903 ∼ 7302 gene families for the studied species to calculate the relationship between gene family size and evolutionary rate. Gene families with large sizes tended to have low Ka/Ks, yet there are a few exceptions. We highlighted the gene families with a size ≥5 and Ka/Ks >1 in red (Fig. 4a). We tested the enriched KEGG and GO terms in these highlighted gene families (Fig. 4b and Supplementary Fig. S4, see the whole list of GO enrichment in Supplementary Table S2). The most highly shared enriched KEGG and GO terms are related to plant-pathogen interactions, signaling, and metabolite biosynthesis.

**Fig. 4.**
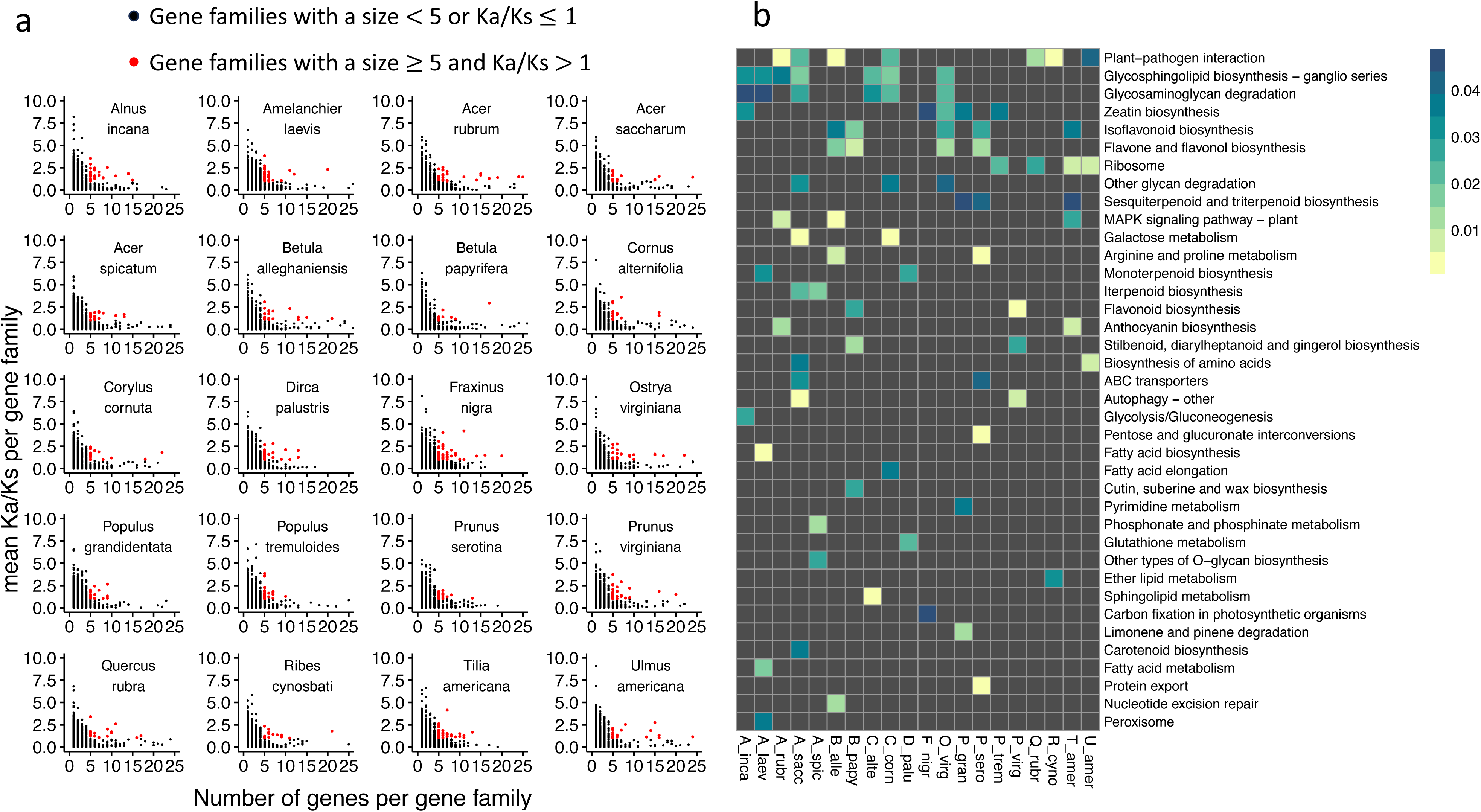
Scatter plot between gene size and Ka/Ks per gene family (a) and KEGG pathway terms enrichment (b) in gene families with a size ≥5 and Ka/Ks >1. The bar in b represents the adjusted p-values of KEGG enrichment test.

### Gene family expansion and contraction

We inferred the birth-death parameter λ (number of gene gains and losses per gene per million years), which is specified in different clades, using the program CAFE5. λ1 (saxifragales) =0.0034, λ2 (rosids) =0.0042, λ3 (asterids) =0.0027, which means that the lineages leading to rosids have the highest gene family turnover rates, followed by saxifragales, and then by asterids. The percentage and numbers of gene family expansion and contraction are shown in Fig. 5a and Supplementary Fig. S5, respectively. All of the studied species have more rapidly expanding than rapidly contracting gene families (family-wide p-value <0.01, Fig. 5b), which may contribute to their lineage-specific novelty.

**Fig. 5.**
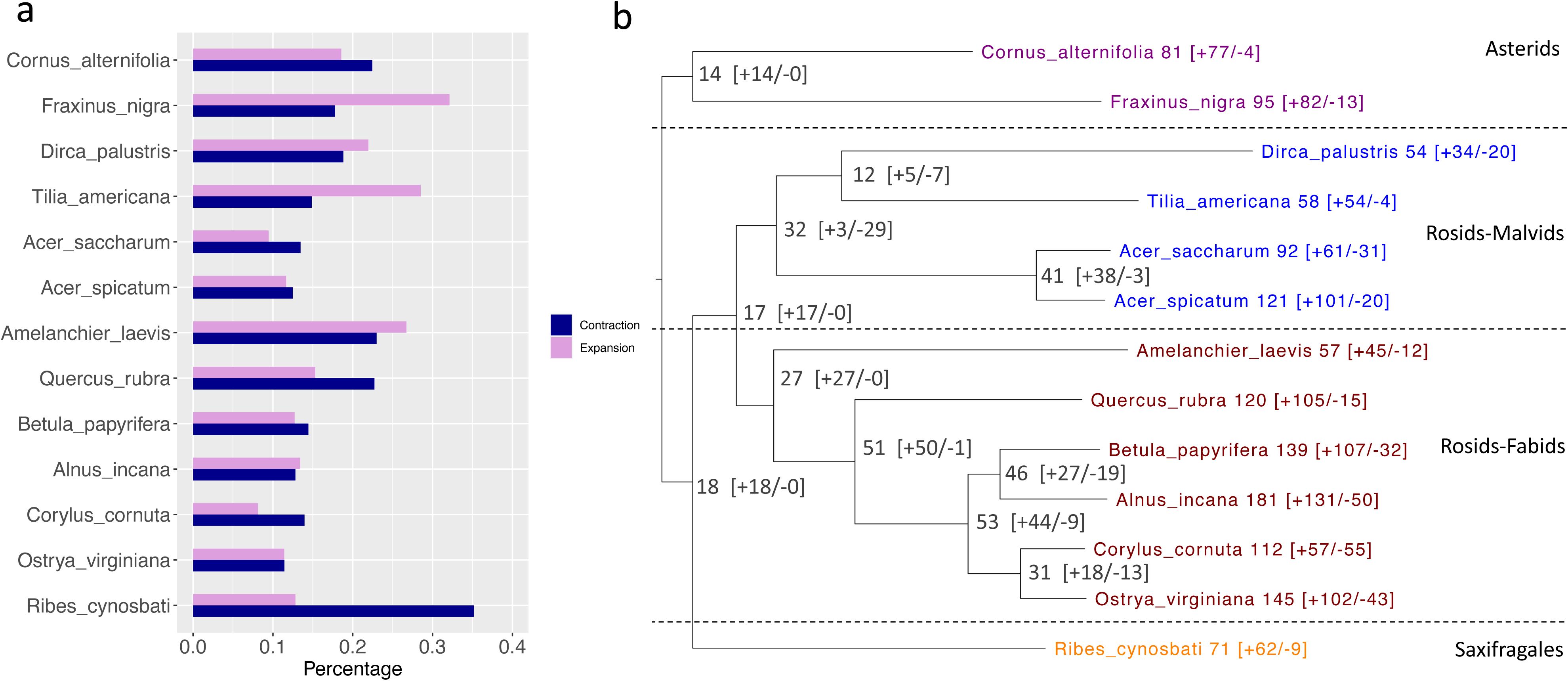
The percentage of gene family expansion and contraction (a) and the number of fast evolving gene families (b), with + representing fast expanding gene family and – representing fast contracting gene family.

We tested the enriched GO and KEGG terms in the fast evolving (expanding or contracting) gene families. All of the studied species shared the enriched KEGG pathway term plant-pathogen interaction (Fig. 6). This term was detected in the fast-expanding gene families of *Cornus alternifolia*, *Tilia americana*, *Acer spicatum*, *Betula papyrifera*, *Quercus rubra*, *Amelanchier laevis*, *Ribes cynosbati*; and this term was detected in the fast-contracting gene families of *Fraxinus nigra*, *Dirca palustris*, *Alnus incana*, *Corylus cornuta*, *Ostrya virginiana*; also, this term was detected in both of the fast expanding and fast contracting gene families of *Acer saccharum*. The gene families assigned to the plant-pathogen interaction term encode proteins (Supplementary Fig. S6) related to disease resistance protein, leucine-rich repeat receptor like protein kinase and others. Some species shared the enriched KEGG pathway terms Tryptophan metabolism and MAPK signaling pathway (Fig. 6).

**Fig. 6.**
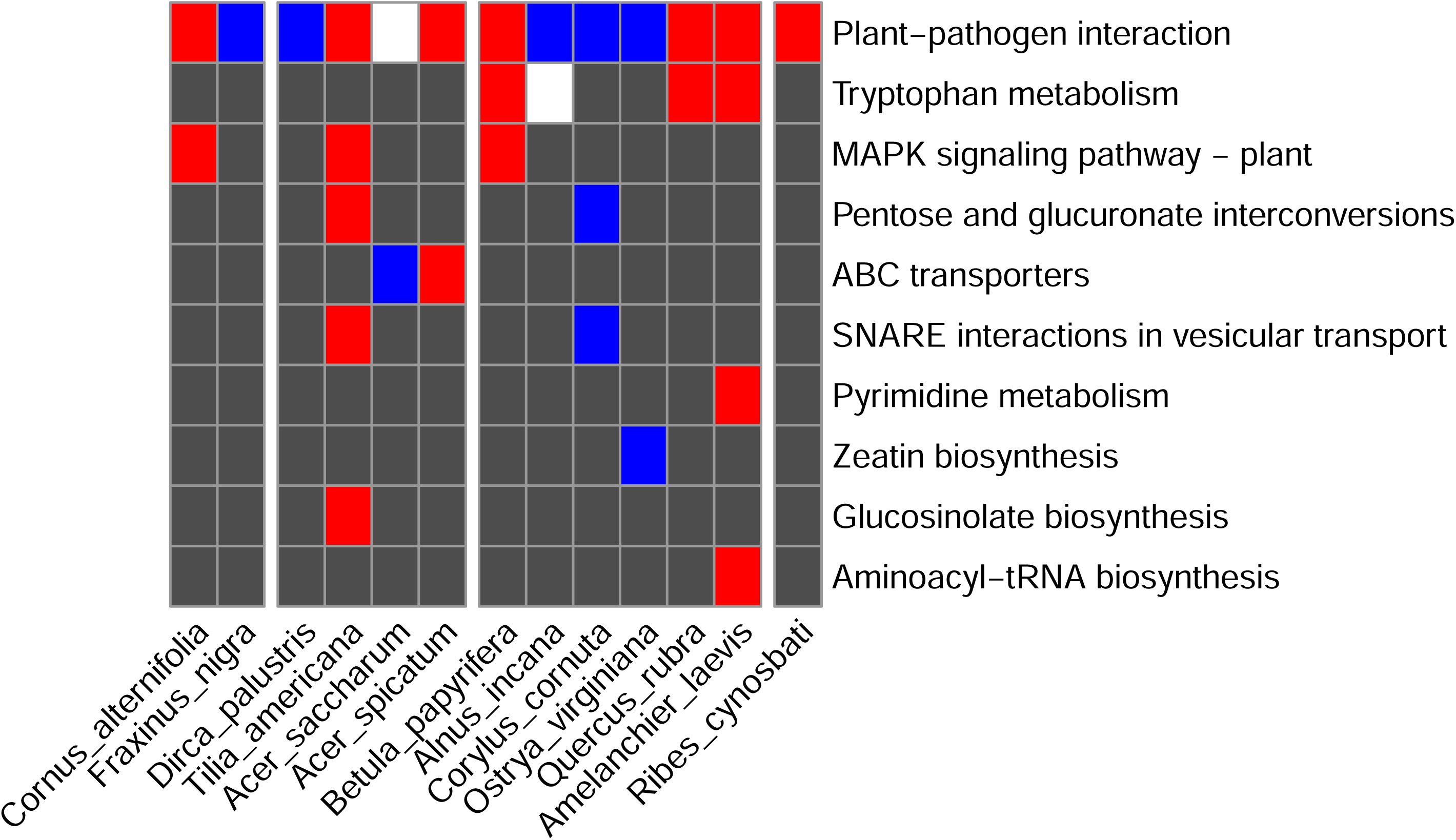
Enriched KEGG pathway terms in the studied diploid species. Red represents fast expanding gene family; blue represents fast contracting gene family; white represents both fast expanding and contracting gene family. Dark grey represents the KEGG terms were not enriched in fast expanding or contracting gene family.

## Discussion

Gene duplication events are prevalent in all organisms and provide genetic variation for evolutionary innovation and adaption to fluctuating conditions (Panchy et al. 2016; Van de Peer et al. 2009; Zhang 2003). Research on the mechanisms of gene duplication as well as the origin, maintenance, and functional roles of genetic diversity in forest tree species are few due to the limited genome and transcriptome data resources. In the present study, we assembled twenty transcriptomes from forest trees that co-occur. We identified the signal of common WGDs shared by core eudicots, and the signal of recent duplication events specific to certain species. We found gene families involved in plant-pathogen interaction, signaling, and metabolite biosynthesis actively remain extensive variation. We also found fast expansion or fast contraction in plant-pathogen interaction genes amongst the studied diploid species. In the following sections, we will discuss these results in detail.

### Tempo and mode of gene duplication

The Ks distribution plots confirmed the widespread paleopolyploidy in plants, that’s said, today’s diploid species might be polyploid (paleopolyploid) at some points in evolutionary history (Blanc and Wolfe 2004). The abundant duplication events provide resources for genetic variation. It is worth noting that Ks distribution is not suitable to infer very ancient events due to saturation with multiple substitutions (Blanc and Wolfe 2004), moreover, different peaks might not be separable from each other, thus it is possible that we missed identifying some duplication events in the present study. Collinearity-or synteny-based methods along with chromosome-level reference genome assembly is an alternative method to identify ancient WGD, yet high quality genome assembly is not easy to obtain (Zwaenepoel and Van de Peer 2019).

The predicted species tree (Fig. 3) generally agrees with current phylogenetic hypotheses for the angiosperms with the exception of the two *Populus* species, which is probably caused by the incomplete gene sets obtained from one leaf RNA-seq sample and/or interspecific gene flow. The duplication events at the root (Fig. 3) of the tree is likely to be related to the gamma WGD before the divergence of rosids and asterids. Such large-scale duplication may contribute to genome structure variation, organismal complexity, and rapid speciation in angiosperm species. WGD events followed by different evolutionary events might also occur in some clades. For *Acer*, it is likely that *Acer spicatum* experienced a diploidization event to revert to diploid status, while *Acer rubrum* maintained a polyploid status.

Divergence time mostly agrees with the previous reports, which stated rosids and asterids diverged about 125 MYA, malvids and fabids diverged about 110 MYA (Zeng et al. 2017). However, divergence estimate in the present study is rather rough. Further studies need to consider using dense taxon sampling, diverse markers like nuclear internal transcribed spacer or chloroplast DNA fragments, and molecular dating with fossil corrections methods to better resolve the evolutionary relationships.

### Sustained large size gene families

Though most duplicate genes were eliminated or pseudonized (Panchy et al. 2016), a small portion of gene families can actively maintain duplicate genes as selection favors the fitness related neo-functionalization. The main functions of these sustained large size gene families are mainly involved in plant-pathogen interaction, signaling, and metabolite biosynthesis (Fig. 4b & Supplementary Fig. S4). This is consistent with previous report that genes encoding functions like signal transduction and stress response tend to have paralogs (Panchy et al. 2016), and genes related to the function like transcription regulation and signal transduction had higher possibility to survive in multiple rounds of genome duplication (Seoighe and Gehring 2004). Plants constantly combat with pathogens and pests. The major class of plant disease resistance (R) genes contain a nucleotide-binding site (NBS) and C-terminal leucin-rich repeats (LRRs). Tandem and segmental gene duplication explain much of the diversification among R genes (Andersen et al. 2020; Leister 2004; Meyers et al. 2003), possibly because they have experienced frequent deletion and duplication throughout evolution, and thus are highly variable (Gao et al. 2018; Hanada et al. 2008). Plant secondary metabolites, such as monoterpenoid, flavonoid, benzoxazinoid, play important roles in defense against predation and pathogens (Bergman et al. 2020; de Bruijn et al. 2018; Falcone Ferreyra et al. 2012). The regulation of plant secondary metabolism requires the intimation of signal transduction network that leads to activation of biosynthetic genes (Zhao et al. 2005).

Gene duplication and deletion give rise to gene family expansion and contraction. The high gene gain rate likely reflects the occurrence of multiple WGDs and frequent SSDs (Carretero-Paulet et al. 2015). While for the other species with contraction outnumbered expansion, the pattern is mainly caused by gene loss and retention of specific gene families followed the common WGDs preceded their common ancestor (Fig. 3). We found a high gene family turnover rate in rosids, but we must caution that the unbalanced taxon sampling in our study species might cause this high turnover value. Moreover, future studies should consider distinguishing the branches with or without recent WGDs, as the branches with recent WGDs tend to have a high gene turnover rate (Casola and Koralewski 2018).

### Rapid evolution in plant-pathogen interaction genes

We found rapid evolution of genes involved in plant-pathogen interaction, tryptophan metabolism and MAPK signaling pathway (Fig. 6) in the adaptive expansion and adaptive contraction gene families across the studied species. The specific expanding or contracting gene families involved in plant-pathogen interaction in each species differed from each other, implying lineage-specificity in changed gene families (Supplementary Fig. S6). This result is consistent with previous studies, which found that fast evolving gene families were mostly groups with functions related with immune defense/stress response, metabolism, cell signaling, chemoreception, and reproduction (Demuth and Hahn 2009). Additionally, a study on *Juglans* (walnuts) found notable contractions in gene families that are involved in disease resistance and other abiotic stress response (Trouern-Trend et al. 2020). Both metabolism and MAPK signaling pathway are associated with plant defense. MAPK cascades sense pathogen/microbe-associated molecular patters and pathogen effectors, and thus play pivotal roles in signaling plant defense against pathogen attack (Meng and Zhang 2013). Plant secondary metabolites contribute to plant immunity as antibiotics, or to control callose deposition and programmed cell death (Piasecka et al. 2015). The biotic stress exerted by pathogen or pests is a major source of selection pressure on the studied trees, which drive the rapid radiation of diverse and large plant immune receptor gene repertoires. Expansion and/or contraction of cell-surface receptor genes correlate with the expansion and/or contraction of intracellular receptor genes (Ngou et al. 2022).

### Limitation of the current study

Our conclusions are drawn from samples collected from co-occurring species from a single forest. The forest type sampled and the species are typical for the entire region including the Ottawa, Nicolet, and Hiawatha National Forests in northern Wisconsin and the Upper Peninsula of Michigan, USA. As individuals within species vary in their adaptive and plastic responses to the environment, our results are, therefore, most applicable to the study region and not the entire geographic range of the species. Furthermore, the assembled transcripts of each species in this study are derived from leaf material and, therefore, cannot represent the whole gene set expressed by the organism. Lastly, we also note that we filtered out genes that had low expression levels to ensure the quality of the assembled transcriptome. Thus, our study does omit some genes, and it is possible that we might end up with different conclusions if we had more genes.

## Conclusions

In summary, we found the common WGDs shared by all the studied species as well as signals of recent duplication events specific to certain species, which are suggestive of different evolutionary patterns. Gene families involved in plant-pathogen interaction, signaling, and metabolite biosynthesis actively remain extensive variation and are fast expanding or contracting amongst the studied species. This knowledge calls for an emphasis of gene family size variation, i.e. copy number variation, when studying forest genetic diversity, which will inform forest adaptive strategy. Future studies may focus on classification of duplications in forest tree species, and investigate how the duplications give rise to species-specific innovation and adaption.

## Data availability

The raw RNA-seq data were deposited in NCBI SRA (accession number: PRJNA896247; http://www.ncbi.nlm.nih.gov/sra). The assembled transcriptomes and their annotation were deposited in Dryad https://doi.org/10.5061/dryad.sj3tx968b. The scripts used for analyses were deposited in https://github.com/Mengmeng-Lu/Gene-duplication-and-gene-expansion-analyses-in-twenty-tree-species.

## Acknowledgements

We thank the Center for Research Computing at University of Notre Dame for high performance computing support. We specially acknowledge the assistance of Dr. Dodi Heryadi, Dr. Scott Hampton, and Júlia Simon. We appreciate Dr. Sam Yeaman and Dr. James R. Whiting for constructive suggestions. The research was performed at the University of Notre Dame Environmental Research Center that has been generously supported by the Bernard J. Hank Family Endowment. The research was permitted under UNDERC research permit number: UNDERC-2021-2.

