## Supplementary Fig.S1-S6 & Table S1 for "Comparative transcriptomics reveals divergence in pathogen response gene families amongst 20 forest tree species"

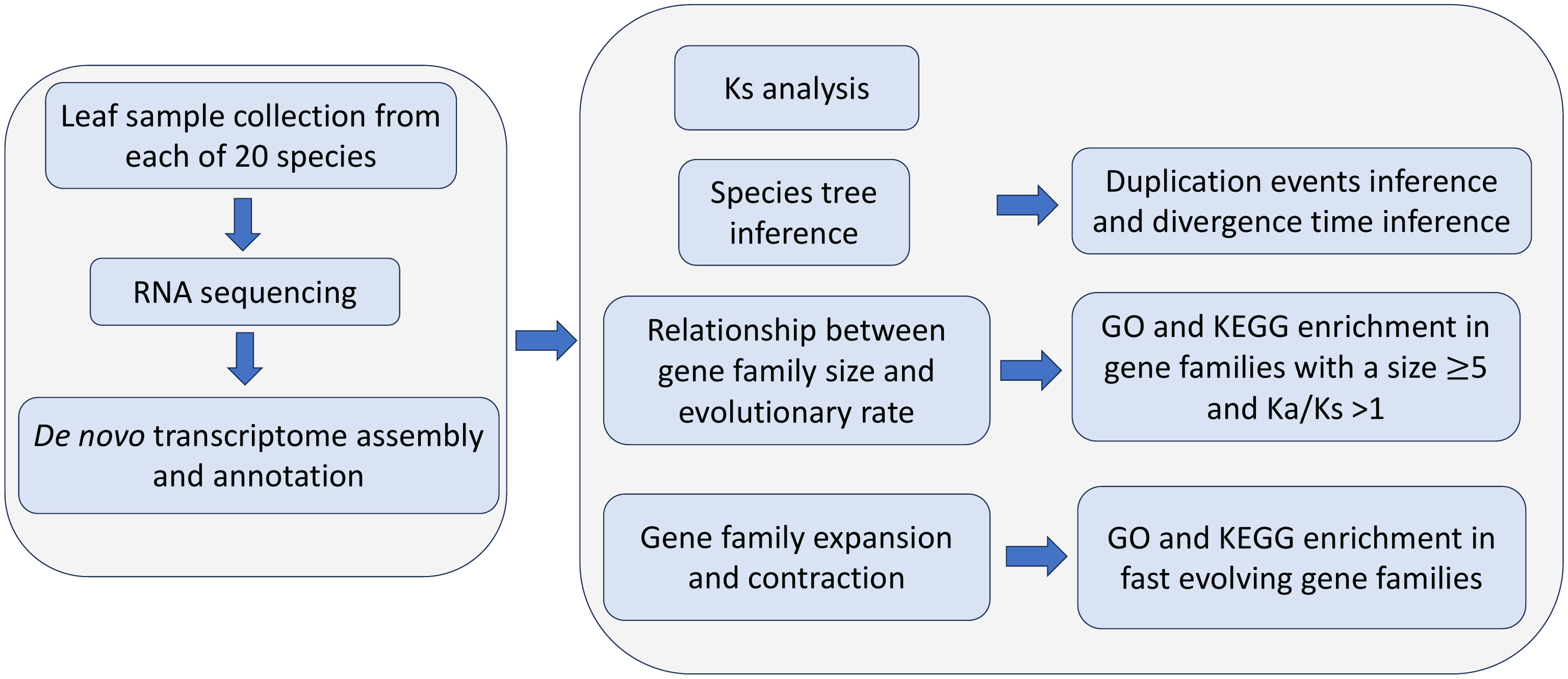

Fig. S1. Research methodology.

Table S1. Quality of the sequence data and the number of assembled transcripts. Raw reads - total amount of reads in raw data; Effective - (Clean reads/Raw reads)*100%; Error - base error rate; Q20, Q30 - (Base count of Phred value >20 or 30) / (Total base count); GC - (G & C base count) / (Total base count)

| Sample | Raw reads | Raw data  (Gb) | Effective  (%) | Error(%) | Q20  (%) | Q30  (%) | GC  (%) | Number of transcripts |
| --- | --- | --- | --- | --- | --- | --- | --- | --- |
| *Acer rubrum* | 42516252 | 6.4 | 99.03 | 0.03 | 96.86 | 91.99 | 43.55 | 26798 |
| *Acer saccharum* | 40258404 | 6 | 99.09 | 0.03 | 96.56 | 91.48 | 44.93 | 19713 |
| *Acer spicatum* | 40951588 | 6.1 | 99.01 | 0.03 | 96.77 | 91.86 | 44.39 | 20529 |
| *Alnus incana* | 40746306 | 6.1 | 98.63 | 0.03 | 97.07 | 92.51 | 46.61 | 25900 |
| *Amelanchier laevis* | 40642302 | 6.1 | 98.61 | 0.03 | 96.81 | 91.95 | 47.28 | 28845 |
| *Betula alleghaniensis* | 40110170 | 6 | 98.77 | 0.03 | 96.75 | 91.86 | 46.77 | 21388 |
| *Betula papyrifera* | 42242750 | 6.3 | 99.14 | 0.03 | 96.87 | 92.18 | 46.57 | 24199 |
| *Cornus alternifolia* | 39167342 | 5.9 | 99.59 | 0.02 | 98.25 | 94.87 | 45 | 22372 |
| *Corylus cornuta* | 42406264 | 6.4 | 99.69 | 0.02 | 98.21 | 94.77 | 46.92 | 19459 |
| *Dirca palustris* | 39318736 | 5.9 | 99.67 | 0.02 | 98.35 | 95.11 | 48.22 | 24726 |
| *Fraxinus nigra* | 40215038 | 6 | 99.54 | 0.02 | 98.36 | 95.05 | 44.35 | 28355 |
| *Ostrya virginiana* | 40860242 | 6.1 | 99.57 | 0.02 | 98.21 | 94.8 | 47.36 | 21072 |
| *Populus grandidentata* | 42223600 | 6.3 | 99.67 | 0.02 | 98.32 | 94.89 | 44.1 | 25882 |
| *Populus tremuloides* | 40203804 | 6 | 99.23 | 0.03 | 96.95 | 92.14 | 44.81 | 25360 |
| *Prunus serotina* | 40248228 | 6 | 99.65 | 0.02 | 98.33 | 94.97 | 45.48 | 24729 |
| *Prunus virginiana* | 40791586 | 6.1 | 98.18 | 0.03 | 96.91 | 92.23 | 46.07 | 24768 |
| *Quercus rubra* | 43706072 | 6.6 | 98.05 | 0.03 | 97.41 | 93.1 | 44.47 | 25300 |
| *Ribes cynosbati* | 40106626 | 6 | 98.97 | 0.03 | 96.57 | 91.52 | 44.57 | 21203 |
| *Tilia americana* | 41811102 | 6.3 | 99.06 | 0.03 | 96.81 | 91.92 | 44.31 | 31804 |
| *Ulmus americana* | 43900232 | 6.6 | 98.69 | 0.03 | 97.7 | 93.78 | 46 | 26486 |

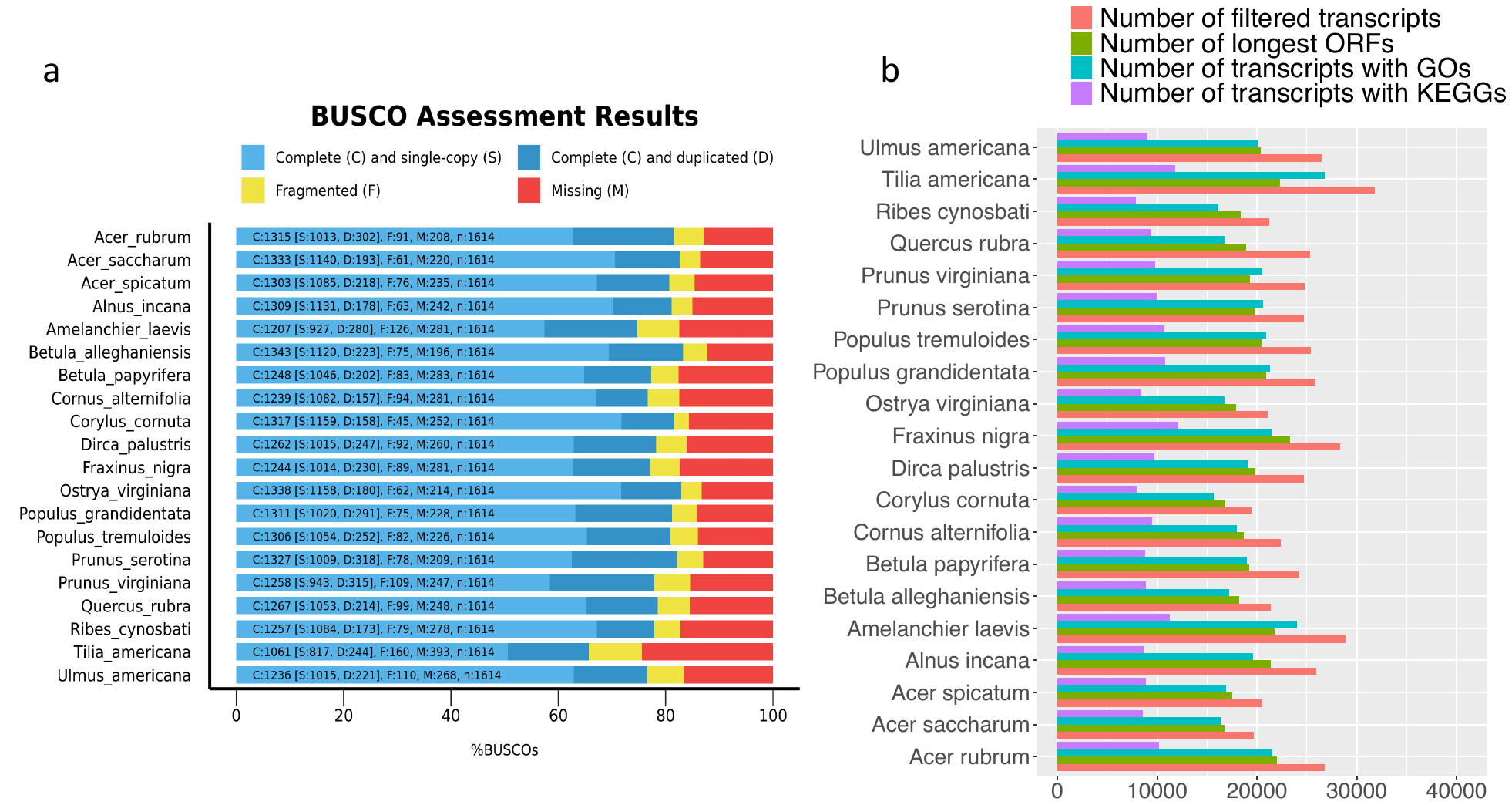

Fig. S2. Completeness assessment of assembled transcriptomes using BUSCO with dataset embryophyta_odb10 (a), and numbers of annotated transcripts (b).

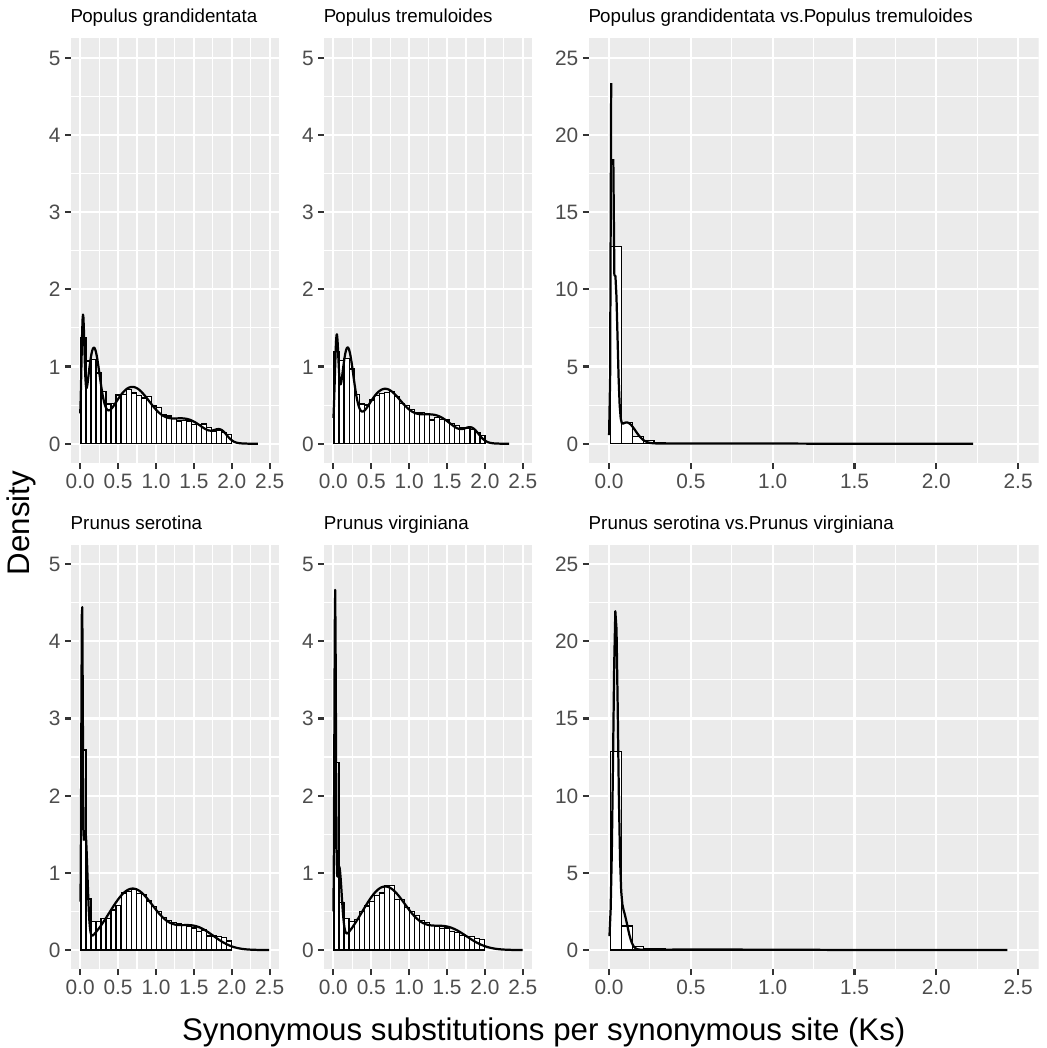

Fig. S3. Distribution of Ks (the level of synonymous substitutions) within and between the *Populus* and *Prunus* species.

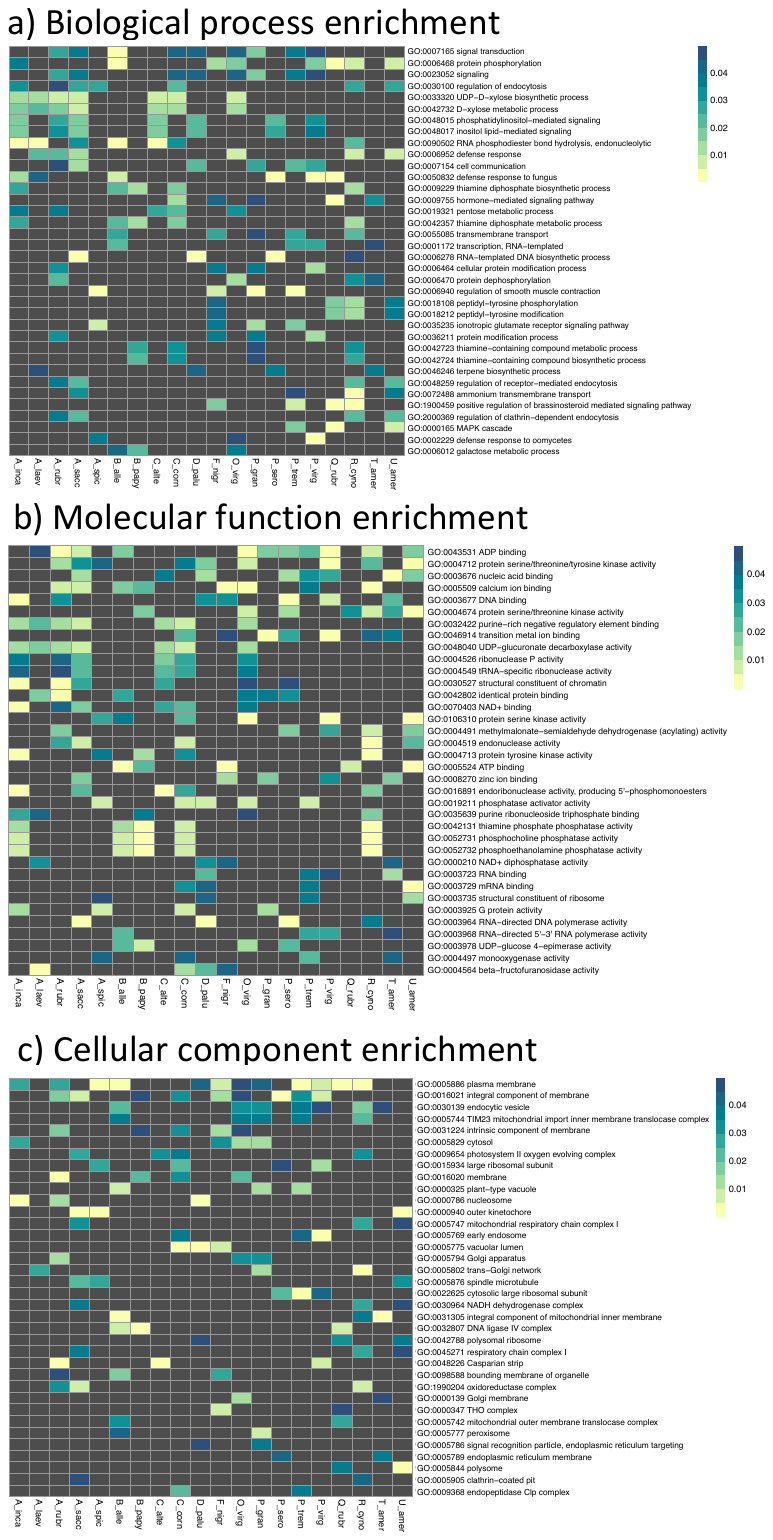

Fig. S4. GO terms enrichment in gene families with a size $\geq$5 and Ka/Ks $>$1. A_rubr *- Acer rubrum*; A_sacc - *Acer saccharum*; A_spic - *Acer spicatum*; A_inca - *Alnus incana*; A_laev - *Amelanchier laevis*; B_alle - *Betula alleghaniensis*; B_papy - *Betula papyrifera*; C_alte - *Cornus alternifolia*; C_corn - *Corylus cornuta*; D_palu - *Dirca palustris*; F_nigr - *Fraxinus nigra*; O_virg - *Ostrya virginiana*; P_gran - *Populus grandidentata*; P_trem - *Populus tremuloides*; P_sero - *Prunus serotina*; P_virg - *Prunus virginiana*; Q_rubr - *Quercus rubra*; R_cyno - *Ribes cynosbati*; T_amer - *Tilia americana*; U_amer - *Ulmus americana*. The color bars represent the *P*-values of GO enrichment test.

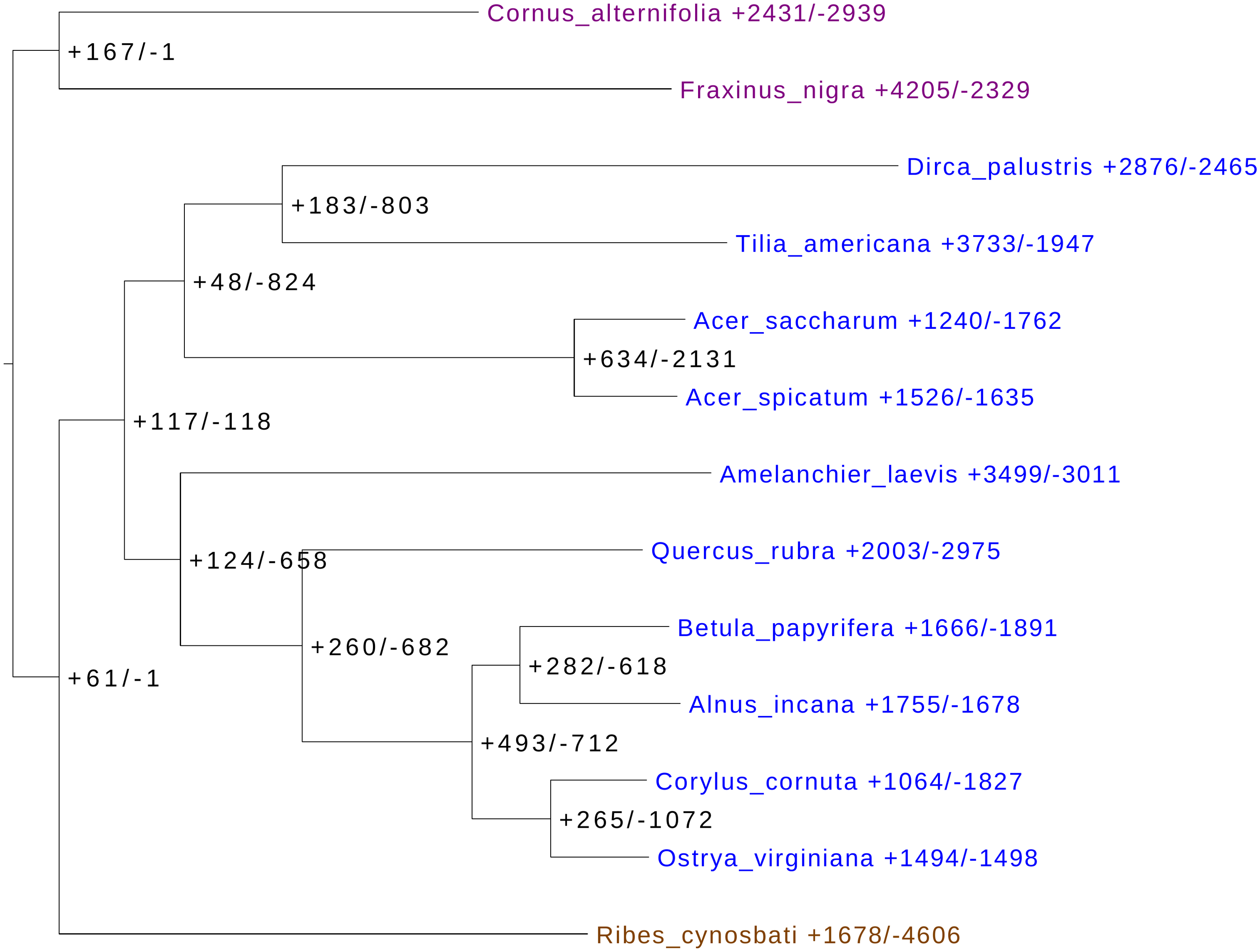

Fig. S5. Numbers of expanded (+) and contracted (-) gene families predicted using the program CAFE5 amongst diploid species.

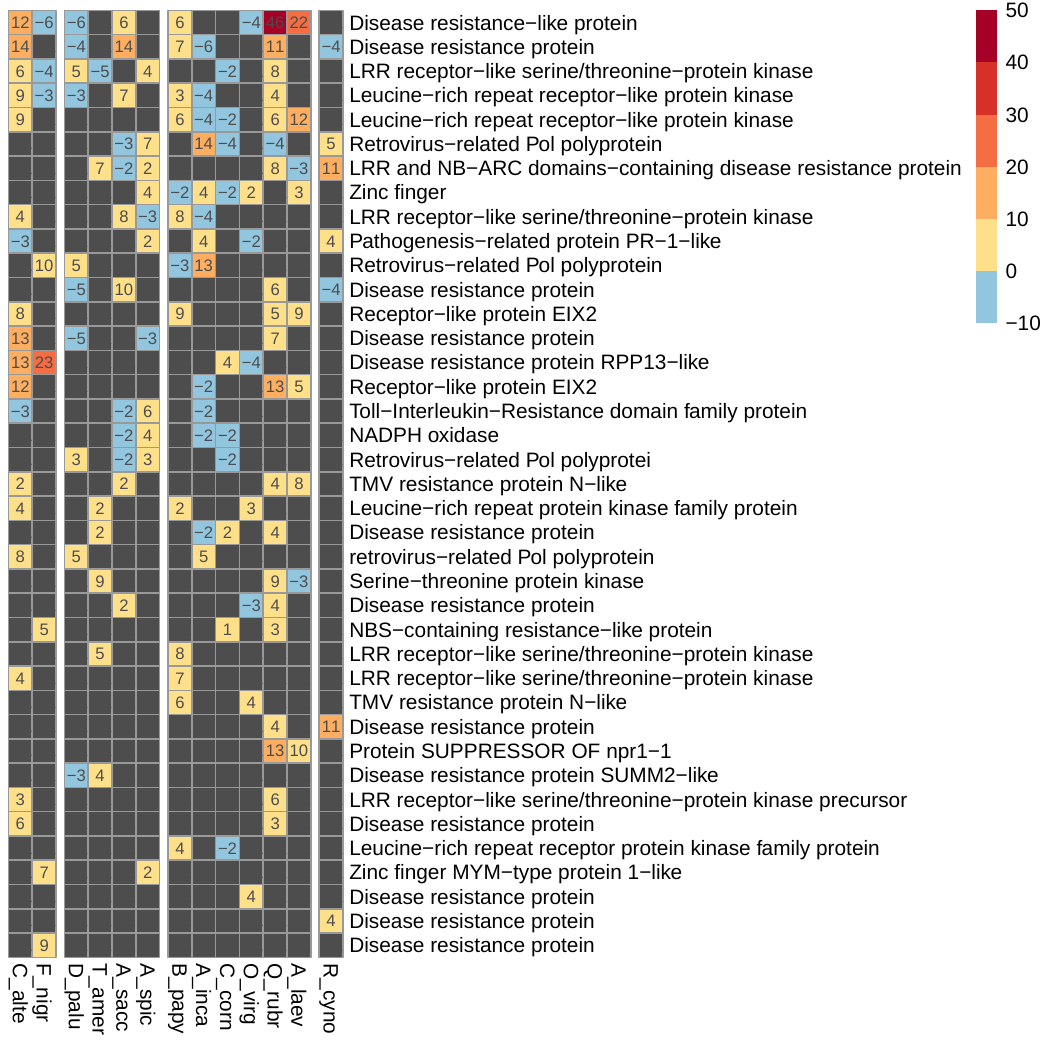

Fig. S6. Numbers of expanded (+) or contracted (-) gene families attached with the KEGG term plant-pathogen interaction. Some gene families may have the same gene annotation. The dark grey cells represent the annotation terms were not found in the expanded or contracted gene families; the color bar represents the number of gene families with expansion (positive value) or contraction (negative values). C_alte - *Cornus alternifolia*; F_nigr - *Fraxinus nigra*; D_palu - *Dirca palustris*; T_amer - *Tilia americana*; A_sacc - *Acer saccharum*; A_spic - *Acer spicatum*; B_papy - *Betula papyrifera*; A_inca - *Alnus incana*; C_corn - *Corylus cornuta*; O_virg - *Ostrya virginiana*; Q_rubr - *Quercus rubra*; A_laev - *Amelanchier laevis*; R_cyno - *Ribes cynosbati.*
